# Dopamine Dynamics in the Nucleus Accumbens Reflect Confidence in Detecting the Occurrence and Non-Occurrence of Visual Signals in Perceptual Decision-Making

**DOI:** 10.1101/2025.04.24.650502

**Authors:** L.J.F. Wilod Versprille, C. McKenzie, F.E. Amorim, K. Yano, J.W. Dalley, T.W. Robbins

## Abstract

Dopamine (DA) is critically involved in processes such as reward anticipation, attention, and decision – making. The present study examined the temporal dynamics of phasic DA transients in the nucleus accumbens core (NAcC) during a visual decisional task based on signal detection theory, using the fluorescent DA sensor dLight1.3b. During the decision-making phase, DA transients in the NAcC encoded real-time outcome expectancy, apparently reflecting the confidence of rats in their choices. Reward prediction errors (RPEs) emerged following reward delivery and omission and were amplified under conditions of increased uncertainty, produced either by degrading the visual target or introducing interfering distraction. Moreover, DA transients were elicited on both visual signal and no-signal trials. These findings demonstrate that DA fluctuations in the NAcC reflect the RPE that incorporates confidence and levels of uncertainty, emphasizing an involvement of nucleus accumbens DA in adaptive decision-making.

## Introduction

Dopamine (DA) is engaged in various behavioural processes including attention, reward, motivation, decision-making, and motor responding ^1,2,11–15,3–10^. DA release occurs in response to predictive cues associated with reward as well as following reward and punishment omission in line with the theory of reward prediction error (RPE), which associates increases in DA with favourable or better-than-expected outcomes and reductions with unfavourable or worse-than-expected outcomes ^16,17,26–29,18–25^. According to Schultz’s RPE hypothesis DA signalling initially occurs at reward presentation but during learning this signal gradually shifts towards cue onset rather than following reward presentation ^16,22,38–47,30–37^. More recently, Gershman and colleagues (2024) proposed a more comprehensive RPE theory, where DA transients reflect RPEs in response to reward while being updated in accordance with the value decay model. Thereby, these transients drive learning, action selection, motivation and vigour ^22,48–51^. DA RPE transients in response to learning under various conditions have been well studied. However, the possible role of striatal DA in well-trained, steady-state performance, for example, decision-making during signal detection tasks has been scarcely investigated, although there is considerable evidence that striatal DA-manipulations affect steady-state performance ^52–59^.

Behavioural paradigms including signal detection tasks, potentially engage a multifactorial DA signalling pattern due to their engagement of simultaneous processes mediated by DA. Disentangling time-dependent DA fluctuations and their specific functions is further complicated by poor spatial and temporal resolution and inherent problems in neurochemical specificity of conventional neurochemical sampling methods ^29,60–68^. Consequently, relatively few studies have attempted to measure quasi real-time DA transients during such performance ^41,69–72^.

The present study investigated time-resolved dopaminergic signalling in the nucleus accumbens core during performance of a visual signal detection task using the fluorescent DA receptor sensor, dLight1.3b. This sensor has successfully been used *in vivo* to investigate DA transients in the freely behaving rodent, although no previous studies have used a signal detection paradigm designed to measure indices of discriminative sensitivity (d’) and bias (β) ^73–77^. Whereas d’ reflects such factors as perceptual accuracy and attention, β indicates motivational tendencies such as perceptual or response bias (for example to report signal presence). Notably, the signal detection paradigm utilizes a true no-signal condition requiring top-down attentional control in order to measure perceptual bias or responsivity through measures of correct rejections as well as hits, misses and false alarms ^78–83^. Varying signal durations were utilised to assess DA transients for increasing levels of physical salience, where the no-signal condition controlled for physical salience. Importantly, to date, there has been no evidence for time-dependent DA signals for instrumental no-signal choices.

Therefore, this instrumental visual detection task necessitates an active choice response to discriminate between signal and no-signal trials, likely engaging higher-order decision-making processes. Subjects must operate with an internal decision-threshold, as described by the signal detection theory, to assess whether perceptual inputs exceed this threshold and to respond accordingly. Moreover, subjects can report on the accuracy of their own decision –making processes, reflecting metacognitive evaluations of choice confidence, which have been reported across species ^84–87^. Typically, higher levels of confidence correlate with greater accuracy and influences both decision – making strategies and perceptual sensitivity (d’). Consequently, some studies demonstrated metacognitive sensitivity (meta-d’), which quantifies the relationship between confidence and performance ^88–90^. Through the effect of choice confidence on decision-making, it may also shape reward anticipation by modulating expectations and adaptive strategies during uncertainty ^91–96^.

Hypothetically, multifaceted DA transients may occur during visual signal detection performance. Specifically, an increase in phasic DA release may occur upon (1) cue presence and omission (on specific no-signal trials); (2) during the decision-making aligned to reflect reward expectancy; (3) reward presentation, with a decrease following non-rewarded incorrect choices.

## Results

### Signal detection performance

Subjects were trained on a visual signal detection paradigm, the signal detection task (SDT) ^97–99^, prior to surgical intervention and subsequent photometry recordings (Fig. 1). The SDT required rats to repeatedly detect and accurately report the presence of a visual cue. This self-paced instrumental choice task was repeatedly performed to obtain a stable-baseline performance. Repeated testing on the SDT yielded stable and reliable behaviour across sessions (see supplementary data); consequently, behavioural and photometry data were averaged over three baseline days. Both the correct detection of the presence of the visual cue (Hit) and its absence (correct rejection (CR)) resulted in a reward. Analysis based on the signal detection theory outcomes (hits, miss, false alarms (FAs), and CRs) revealed significant differences in choice outcome (Fig. 2A; F_(3, 56)_ = 87.445, p < 0.001) and choice latency (Fig. 2C; F_(3, 162)_ = 6.93, p < 0.001). There was a general bias to report the signal (mean ± SEM: β = 0.84±0.06), while exhibiting an average sensitivity of d’ = 1.00±0.13. The attentional load of the task was enhanced by varying the signal duration of the visual cue (30ms, 60ms, 250ms and 1s), with physically more salient signals resulting in higher accuracy compared with shorter signal durations and no-signal trials (F_(4, 56)_ = 26.35, p < 0.001) (see supplementary data).

**Figure 1:**
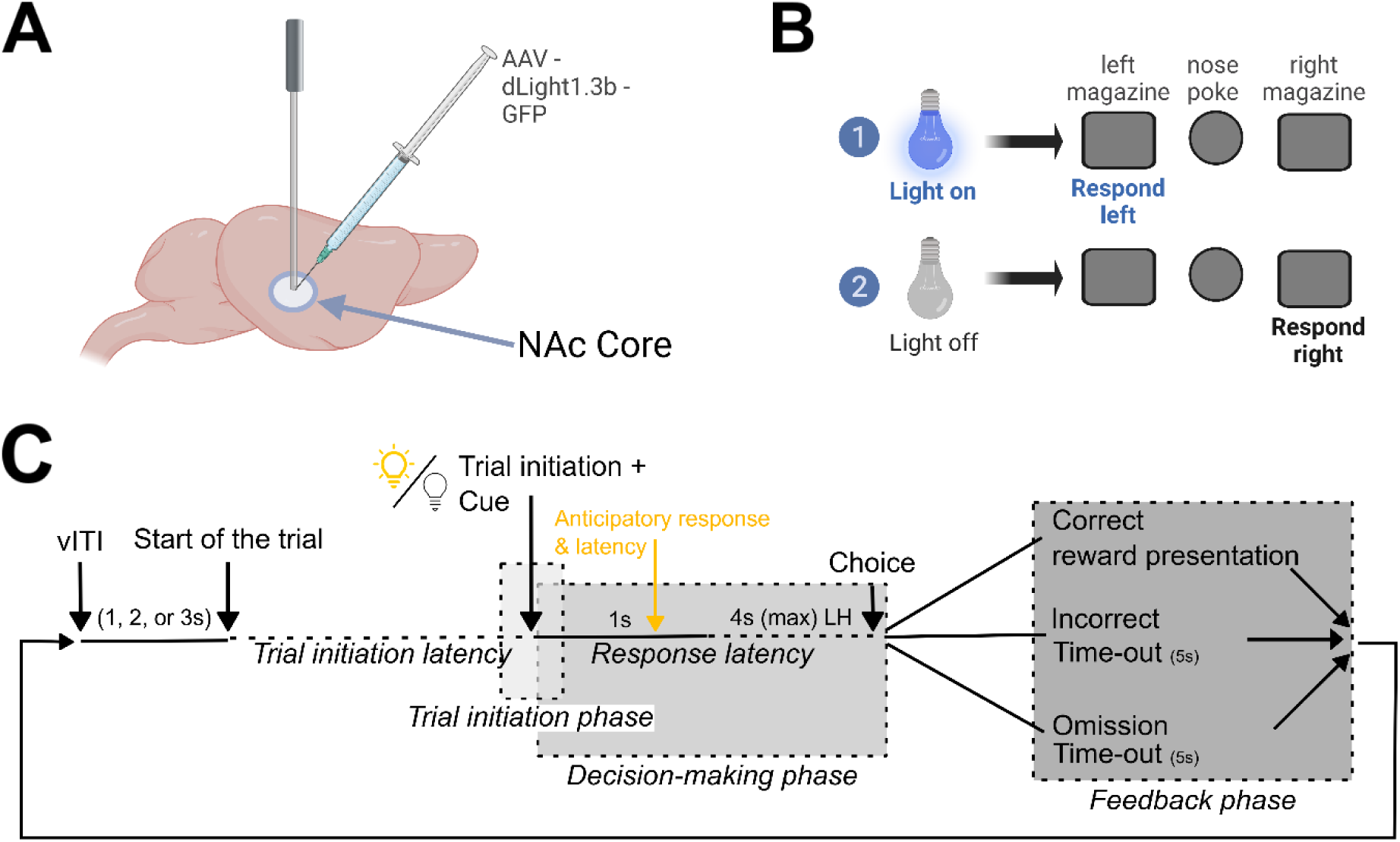
A schematic overview of the experimental procedures. (A) DA signals were recorded following surgical optic fibre implantation and viral infusion of adeno associated vector and promotor of dLight1.3b in the nucleus accumbens core. (B) Subjects were trained on the SDT to correctly identify the presence or absence of a visual cue that was triggered by a nose-poke in the central aperture. (C) Timeline of a typical trial in the SDT. The SDT trials were self-paced; at the start of the trial the light inside the central nose-poke aperture was illuminated indicating that the subject could initiate a trial through a nose-poke, and the trial initiation latency was recorded. The trial initiation immediately prompted the visual cue directly above the centre nose-poke aperture to illuminate or remain off. The interval immediately preceding trial initiation until just after trial initiation was defined as the trial initiation phase. Trial initiation started the decision-making phase, where subjects responded to the respective food magazine (left or right) associated with signal or signal omission. A choice was only recorded after the cue phase (1 s) had elapsed (the response latency was ≥ 1 s), responses made before the end of the 1 s were classified as anticipatory responses with respective choice latency. Following this 1 s period, a choice could be made within the 4 s limited hold (LH) or an omission was recorded and a time-out (onset of the house-light for 5 s) commenced. The choice started the feedback phase, where all correct choices were immediately rewarded, and all incorrect choices were immediately punished with a time-out. Subsequently, there was a variable inter-trial interval (vITI) of 1, 2 or 3 s before the next trial would begin. Abbreviations: LH = limited hold, NAc = nucleus accumbens, vITI = variable Inter Trial Interval. Figure was created in BioRender; Wilod Versprille, L. (2025) https://BioRender.com/b2oe3yx.

**Figure 2:**
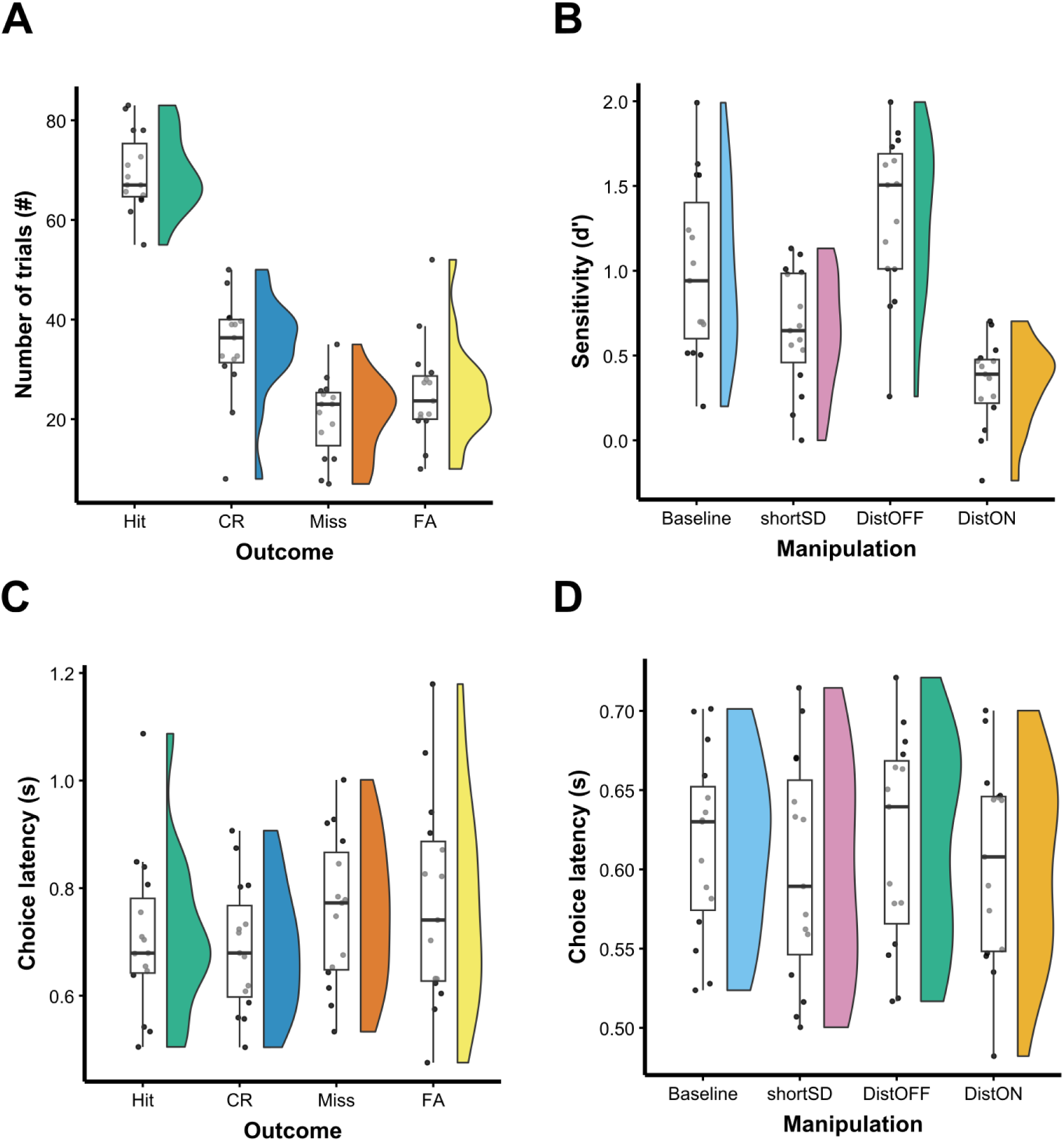
Signal detection performance measures (n = 15). (A) Average number of signal detection theory outcomes across three baseline sessions by subject. Outcomes include hits, correct rejections (CR), miss and false alarms (FA). Mixed linear effects model of signal detection theory outcomes: F_(3,56)_ = 87.4, p < 0.001. Tukey post-hoc tests show significant differences between all outcomes with the exception of misses compared with FA. (B) Sensitivity (d’) across test conditions, where baseline is averaged for each subject across three sessions, shortSD the sensitivity during the vSD manipulation, DistOFF the baseline trials of the intra-distractor manipulation and DistON the interspersed 20 trial blocks with visual distractor. Mixed linear effects model of d’: F_(3,42)_ = 32.38, p < 0.001. Tukey post-hoc tests revealed significant differences between all groups: Baseline – ShortSD (t = 3.2, p = 0.01), Baseline – DistOFF (t = –3.1, p = 0.02), Baseline – DistON (t = 6.2, p < 0.001), ShortSD – DistOFF (t = –6.3, p < 0.001), ShortSD – DistON (t = 3.0, p = 0.02), DistOFF – DistON (t = 9.3, p < 0.001). (C) Average choice latency for each signal detection theory outcome category across three baseline sessions by subject. Mixed linear effects model of signal detection theory outcomes: F_(3, 162)_ = 6.93, p < 0.001. Tukey post-hoc tests revealed significant differences between choice latency of Hits – FA (t = –3.0, p = 0.02), CR – Miss (t = –3.2, p = 0.008), and CR – FA (t = –3.9, p < 0.001). (D) Lack of effect on choice latency across test conditions depicted in (B).

Behavioural manipulations of attentional load (i.e. by either reducing the physical salience of the visual targets through reducing their duration or introducing irrelevant visual stimuli) systematically altered stimulus uncertainty, necessitating greater attentional control (Fig. 2B and D). Two different manipulations were employed to investigate the influence of uncertainty: the short vSD manipulation introduced a novel, rapid signal duration, while the intra-distractor manipulation interspersed blocks of 20 trials with a visual distractor. The manipulations significantly impaired task performance, as evidenced by a reduction in discriminative sensitivity (d’: F_(3,42)_ = 32.38, p < 0.001), but importantly did not affect response bias (β) or response speed (choice latency).

### Fibre photometry recordings during visual signal detection performance

To examine DA dynamics during the visual signal detection task, we transfected dLight1.3b into the nucleus accumbens core (NAcC) of male Sprague Dawley rats. Three weeks later, dLight1.3b functional expression was investigated by recording DA transients to the non-contingent presentation of sucrose pellets. Subsequently, DA transients were repeatedly recorded during the performance of the visual detection task. There was a high consistency of DA transients across trials, days and subjects. Subjects were counterbalanced for signal associated side and hemisphere, and there was no relationship between DA signal and the recorded hemisphere (left or right), the magazine location (left or right) associated with responding signal presence, or the hemisphere (ipsilateral or contralateral) connected to the magazine location associated with signal presence (see supplementary data). Further analysis of the signal into frequency bands revealed that DA activity in the NAcC exhibited slow dynamics (0.15 – 2.5Hz) as previously reported by Jørgensen and colleagues (2023) (see supplementary data) ^100^.

### DA transients during signal detection task performance resemble RPE-like signals

DA-transients were measured individually in well-trained rats over the course of three consecutive baseline SDT sessions and averaged by subject according to trial type (Fig. 3). The omission of a response in either food magazine within the 4s limited hold were so rare that they were excluded from the analysis. A mixed effects model examining differences in AUC by outcome and time frame revealed significant main effects of outcome (F_(3, 264)_ = 53.38, p < 0.001) and time frame (F_(5, 264)_ = 29.44, p < 0.001) as well as an outcome x time frame interaction (F_(15, 264)_ = 12.77, p < 0.001). DA consistently spiked at trial-initiation irrespective of choice and reward outcome (Fig. 3A-B). DA elevation centred around the initiation of the trial and discriminative stimulus is the instrumental equivalent of a reward predictive DA signal of a conditioned stimulus. In the feedback phase, DA remained elevated following a correct choice whilst DA levels decreased below baseline DA levels following an incorrect (unrewarded) response (Fig. 3A-B). Notably, in the decision-making phase, DA-transients were outcome dependent, DA remaining elevated in anticipation of a correct choice, whether correctly reporting a visual target or correctly reporting the absence of the target (i.e. correct rejection). By contrast, in anticipation of an incorrect choice, DA-levels began to decrease prior to feedback, again for both signal detection and correct rejections (Fig. 3A-B). DA transients thus behave in a manner consistent with RPE-theory by reflecting a mismatch between expectation and outcome.

**Figure 3:**
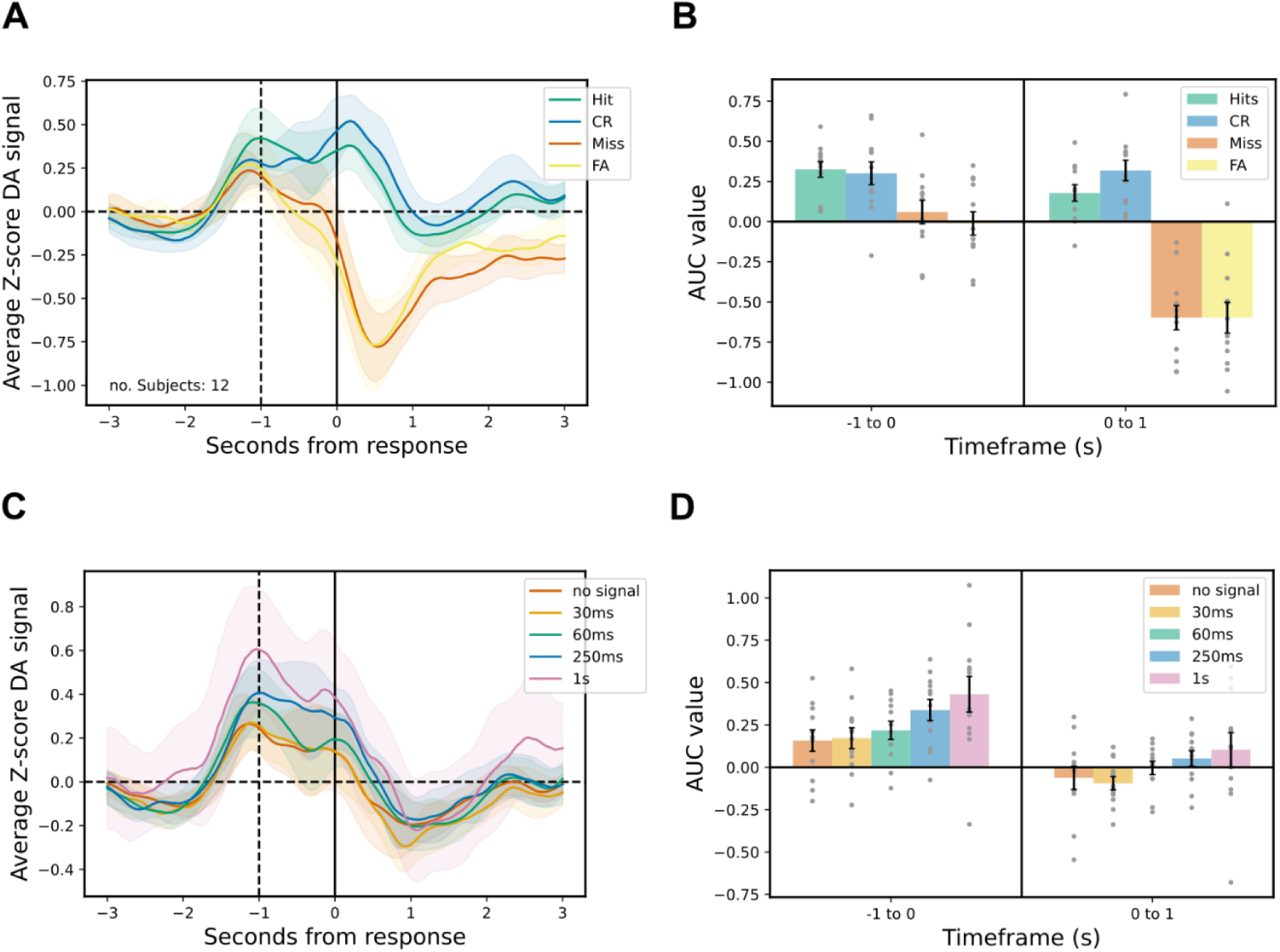
DA transients measured during SDT performance. The left panels (A, C) depict the average Z-score ± 95% confidence interval for each category and the right panels (B, D) depict the area under the curve (mean ± SEM) for every category, expressed in 1s timeframes. Fibre photometry (FP) results were averaged across three sessions. Results were aligned to choice and feedback. (A) DA transients by response type; (B) AUC by response type of –1 to 0 and 0 to 1s timeframes; (C) DA transients by signal duration; (D) AUC by signal duration of –1 to 0 and 0 to 1s timeframes.

There were no significant differences in DA transients for signal trials compared to no-signal trials. Instead, DA transients were affected by outcome, with Hits and CRs as well as Misses and FAs exhibiting similar DA signalling profiles. These findings indicate that the absence of a cue produces a discriminative response, reward predictive signal, and RPE signal analogous to that associated with the actual presence of a visual cue.

Signal cues of varying durations were presented, with longer cues (1s, 250 ms) being more physically salient than shorter cues (30ms, 60ms), alongside the no-signal trials without physical salience. The ability to distinguish between short signal cues and no-signal trials differentiated low and high-attentive subjects. Accuracy was higher for longer signal durations, with rats correctly identifying these signals more frequently. This increased certainty associated with longer signal durations corresponded to elevated DA transients (Fig. 3C-D), both at trial initiation and throughout the decision-making phase (main effect of signal duration (F_(4, 319)_ = 6.81, p < 0.001) and time frame (F_(5, 319)_ = 29.79, p < 0.001). Notably, differences in DA amplitude at trial initiation persisted across correct and incorrect outcomes, while rewarded no-signal trials showed a significant increase in AUC during the feedback phase compared to rewarded 60ms and 250ms trials.

### DA transients and Reward anticipation

Outcome-dependent differences in DA transients suggest that reward anticipation is dynamically updated throughout the decision-making process, influencing the RPE in real-time. To investigate this possibility, DA transients of fast (the 25% trials with the most rapid choice latency) and slow responses (the 25% trials with the slowest choice latency) were compared, under the assumption that rapid responses reflect heightened decision-confidence and increased reward anticipation in accordance with Stolyarova and colleagues (2019) (Fig. 4) ^101–104^. Response speed was found to influence DA transients in an outcome-dependent manner (outcome x choice latency: F_(3, 511.38)_ = 2.32, p = 0.07). Thus, rapid correct responses were associated with increased DA release at trial initiation and throughout the decision-making phase compared with slower correct responses (time frame x choice latency: F_(5, 510.87)_ = 1.94, p = 0.09). During the feedback phase, slow CRs exhibited elevated DA transients compared with rapid CRs, rapid Hits, and slow Hits, indicating an increased magnitude of the RPE.

**Figure 4:**
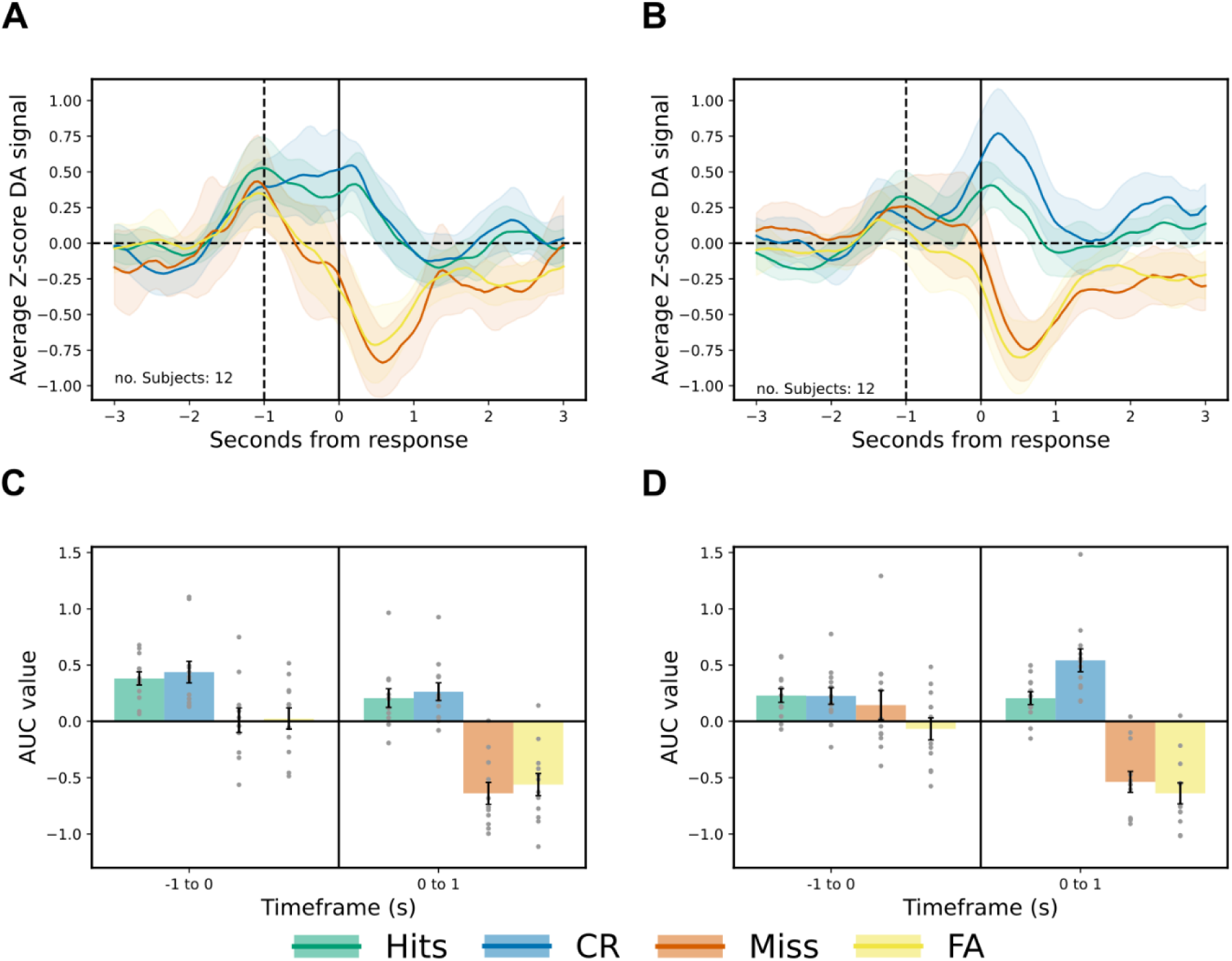
DA transients grouped according to fast and slow anticipatory response times on the. (A) averaged DA transient by response type for the quartile with most rapid choice latency; (B) averaged DA transient by response type for the quartile with the slowest choice latencies; (C) AUC by response type of –1 to 0 and 0 to 1s timeframes for the rapid choices; (D) AUC by response type of –1 to 0 and 0 to 1s timeframes for the slowed choices. FP results were an average of three sessions and depicted as z-score ± 95% confidence interval. Results were aligned to choice and feedback.

### Uncertainty affects DA transients consistent with reward anticipation

We used two behavioural interventions to introduce uncertainty into the task and thereby an increased demand for attentional control (Fig. 5; time frame x behavioural manipulation: F_(15, 1045)_ = 1.69, p = 0.048). The short vSD manipulation did not affect the amplitude of the DA transients at the trial initiation, but did affect DA transients throughout the decision-making phase, where DA transients decreased prior to receiving positive and negative feedback (Fig. 5A-B). Therefore, the outcome-dependent effect seen in the baseline SDT disappeared following this behavioural manipulation. In contrast, the DA transients significantly increased in magnitude following positive feedback and reward presentation, supported by a significant interaction between outcome x time frame (F_(15, 253)_ = 12.72, p < 0.001).

**Figure 5:**
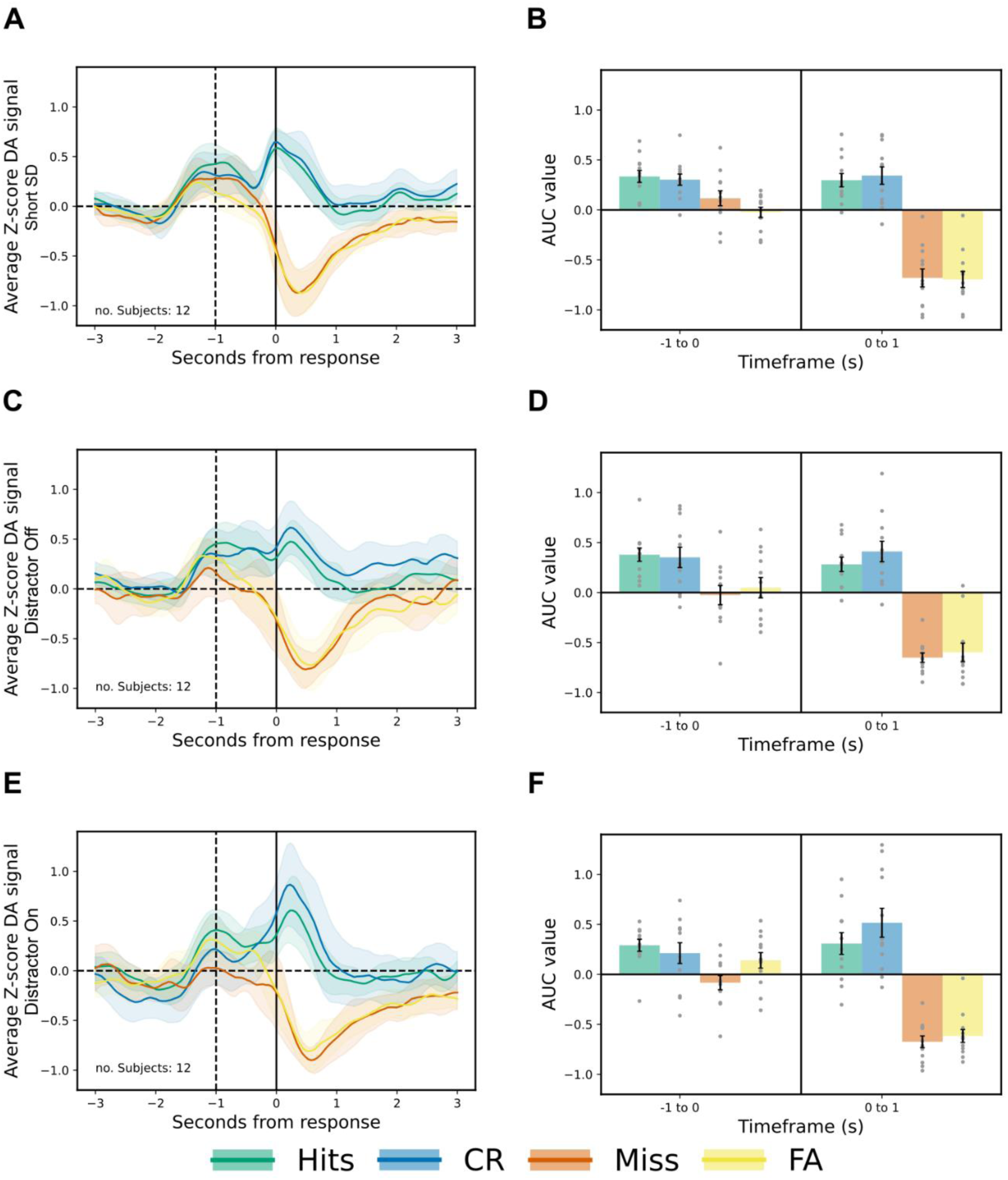
Summary of the effects of increased uncertainty on DA transients during SDT performance. The left panels (A, C, E) depict the average Z-score ± 95% confidence interval for each category and the right panels (B, D, F) have the area under the curve (mean ± SEM) for every category in the –1 to 0 and 0 to 1s timeframes. Results were aligned to choice and feedback. (A & B): These panels represent the DA transients following the short vSD manipulation, with 230 trials instead of 170 due to the addition of 30 further no-signal trials and the addition of very rapid signal durations (30×10 ms). (C – F) Represent the DA transients during the intra-distractor manipulation, with (C & D) depicting the results during the standard SDT trials, and (E & F) depicting the results during the interleaved blocks of trials with visual distractor (house light blinking at 1 Hz from the start of the trial until choice/feedback for blocks of 20 consecutive trials).

The introduction of a visual distractor affected DA transients in similar fashion, both during distraction and in the interleaved standard SDT trials. In accordance with observations made during the short vSD manipulation, the amplitude of the DA transients increased during positive feedback for both standard and distraction trials (Fig. 5C-F; outcome x time frame: F_(15, 517)_ = 16.58, p < 0.001; time frame x distractor: F_(5, 517)_ = 2.43, p = 0.034). For Correct Rejection trials during distraction, there was diminished DA elevation during the decision-making phase, followed by an enhanced amplitude during the feedback phase, indicative of a RPE. Notably, this DA signalling pattern was specific to the Correct Rejection trials during distraction and was not observed for Hits during distraction or non-distraction trials. Moreover, during the trial-initiation phase for both the standard and distraction trials, the DA amplitude for Miss trials was either less pronounced or disappeared entirely, respectively. Thus, reward prediction and errors in reward prediction were dynamically affected by the level of attentional load irrespective of the way it was introduced.

## Discussion

Our findings demonstrate that DA transients in the NAcC during performance of a well-trained visual signal detection task in its decisional phase represent real-time outcome expectancies, whether of success or failure. These transients could be considered to reflect the animals’ confidence concerning the outcome, an important component of decision-making ^51,84,88,105,106^, as more rapid hits and correct rejections were associated with greater amplitude DA transients than slower responses. In this well-trained task, RPEs following reward presentation or absence were still evident. However, when task manipulations heightened volatility and increased attentional load—either by shortening the visual signal duration, thereby increasing uncertainty, or by introducing a visual distractor that interfered with signal processing—DA transients during the decision phase became less differentiated. Instead, these manipulations amplified the magnitude of RPEs produced by reward feedback. A further notable discovery was that DA transients were elicited equivalently on visual signal trials and on those trials where no-signal occurred. There was thus no bias in DA transients caused by visual stimulation and this suggests that the DA responses are most closely associated with the response contingencies associated with these distinct cues rather than their physical salience *per se*.

### DA transients in response to choice-feedback reflect a reward prediction error

As hypothesised, the gradual ramping of DA during the discriminative stimulus in this instrumental choice paradigm appears comparable to DA elevations in response to Pavlovian conditioned stimuli ^16,22,38–47,30–37^. Moreover, DA transients occurring in the feedback phase for correct and incorrect responses were consistent with positive and negative RPEs, respectively. Thus, DA transients in this visual detection task are broadly consistent with Schultz’s RPE-theory, despite occurring during a well-trained self-paced choice paradigm. However, the DA transients observed here more closely followed the RPE two-component response proposed by Schultz (2016), where DA transients act as a utility prediction error signal used for learning, utility and economic decision-making ^22,49,50^. Additionally, for the task manipulations that heightened volatility and enhanced attentional load, DA transients during the decision-making phase were markedly attenuated. However, those following reward in the feedback phase exhibited greater relative AUC and peak size as compared with no reward trials. Thus, the enhanced uncertainty may well represent a novel learning environment which depends on RPE ^107^. The elevated DA seen in rewarded, uncertain trials, consequent upon the ‘surprise’ of a better-than-expected outcome, further supports the utility of the DA transients in learning and also in decision – making as steady-state performance is re-established. In line with these findings, Li and colleagues (2023) also reported that dynamic and rapid inter– and intra-trial feedback influenced neural reward processing ^23^.

### Comparable DA transients on signal and no-signal trials

The absence of the visual cue elicited a DA signalling pattern equivalent to that following its presence. Previous studies have demonstrated that DA transients typically occur in response to exteroceptive cues predictive of reward, particularly in Pavlovian conditioning tasks ^14,24,30,108–111^. These findings are generally interpreted as reflecting reward expectancy. However, the present instrumental visual detection task necessitates an active choice response to discriminate between signal and no-signal trials, which may engage higher-order decision-making processes.

Our findings reveal that elevated DA transients were associated with decision-making for both Hits and Correct Rejections, i.e. the detection of the *absence* of the visual signal, while decreasing for Misses and False Alarms, presumably representing attentional failures. Intriguingly, in Correct Rejection trials, the elevated DA transients were triggered by an interoceptive discriminative stimulus rather than merely occurring simply in reaction to an exteroceptive (visual) cue. This suggests that DA activity arises when a detection threshold fails to be met, highlighting its role in accurately reflecting reward anticipation rather than simply signalling the presence of a sensory cue.

Notably, despite DA transients indicating anticipation of an incorrect choice and subsequent reward omission, subjects did not adjust their decisions. Although this behaviour might suggest a dissociation between the neurochemical representation of reward prediction and behavioural adaptation, it could also be explained by the limited, sub-second time available for subjects to monitor and hence modify their choices. While some studies report similar DA signalling patterns in response to both rewarding and aversive cues ^48,95,96,112^, no prior work has described this pattern in the context of cue non-occurrence requiring an active choice.

### DA transients driving decision-making reflect real-time reward anticipation and confidence

Especially striking was that DA transients during the decision-making phase were modulated by the *subsequent* trial outcomes. Trials with correct outcomes exhibited significantly greater DA elevation compared to incorrect trials, particularly during rapid responses. Clinical studies consistently demonstrate that elevated decision-confidence is commonly associated with such rapid response times, as demonstrated with the confidence database; a collection of 145 datasets with data from over 8,700 participants ^101,103,113^. In contrast, no DA elevation was observed on trials with slowed choice latencies, frequently associated with uncertainty and guessing. This suggests that DA transients during decision-making reflect more than classical RPE signalling; they encode a metacognitive belief in choice accuracy or decision-confidence and engage DA-dependent processes necessary for effective discrimination.

Several studies support the involvement of metacognitive confidence in decision-making in non-human primates and rodents ^84,91,105,114–119^. More precisely, Lak and colleagues (2017) demonstrated that midbrain DA neurons in macaques reflect both reward size and confidence in decision accuracy, aligning with Bayesian decision-making theory ^91^. In a related study, Lak and colleagues (2020) showed that mouse choices integrated sensory confidence, past rewards and learned value, creating a model that bridges signal detection and reinforcement learning principles ^114^. Similarly, Kutlu and colleagues (2021) reported that NAcC DA release in mice reflects the “perceived salience” of stimuli in a discriminative task ^96^. However, in the present study, NAcC DA release was enhanced in anticipation of a correct no-signal detection which presumably is devoid of physical salience.

Intriguingly, DA transients for rapid False Alarm and Miss trials (both involving active responses) were inconsistent with enhanced reward anticipation, instead declining rapidly during the decision –making phase. These rapid, incorrect responses could reflect impulsive decisions made prematurely before visual processing has concluded; hence, an uninformed guess becomes an informed mistake. Alternatively, rapid incorrect responses may not reflect enhanced confidence, as suggested by Overhoff and colleagues (2022), who demonstrated a relationship between response speed and self-reported confidence for correct but not for incorrect choices ^103^.

### Neural circuitry engaged in confidence

Confidence is thought to arise from neural computations integrating sensory input, prior knowledge, and reward prediction. These processes are generally regulated by regions within the frontal cortex such as the medial prefrontal cortex (Lak et al. 2020) or the orbital frontal cortex (Kepecs et al. 2008) ^105,114^. These prefrontal regions affect subcortical regions including the NAcC through top-down signalling pathways. While the precise role of accumbal DA in confidence remains unclear, it is often expected that such signalling would influence action or choice. However, evidence from Vaghi and colleagues (2017) suggests that confidence can sometimes operate independently of immediate action^120^.

The observed NAc DA signals may serve as mechanism that continuously monitors learned, steady-state performance even at asymptote. Hereby, DA signals dynamically evaluate whether behaviour should be flexibly updated in response to alterations in reward expectation as well as reward feedback. Rather than solely guide immediate action, this signalling may modulate longer-term decision-strategies or encode belief states that inform future behaviour as proposed by Lak and colleagues (2017). The frontal cortex likely plays a critical role in integrating signals projecting from the NAcC, as well as possibly other DA-dependent striatal regions such as the NAc shell and dorsal striatum and modifying behaviour appropriately. How cortical regions involved in metacognition influence DA signals in the NAcC requires clarification. Specifically, future studies should investigate whether these signals integrate confidence with learned expectations and executive processes such as behavioural flexibility, task-switching and attention. Moreover, these studies could address the distinct contributions of subcortical regions and the role of other neuromodulators, such as acetylcholine and noradrenaline ^5,7,9,10,121–125^.

### Limitations and further considerations

A notable limitation of this study is its exclusive focus on dynamic changes in phasic DA, driven by the need to relate fast-acting DA transients to rapid behavioural events. In contrast, tonic DA changes, which occur on a slower timescale, were not evaluated due to the inherent challenges in linking these gradual fluctuations to the fast-paced nature of the task. Future studies could explore how tonic DA influences long-term behavioural adaptations and interacts with phasic DA signalling. Additionally, there was some inconsistency in viral transfection, resulting in individual differences in SNR of DA transients, increasing variability in the AUC and amplitude scores despite the use of normalized z-scores. Therefore, caution must be taken when comparing subgroups, such as the hemisphere implanted with the optic fibre, the side associated with the cue, and its combination.

Fibre photometry measures fluctuations in DA concentration relative to baseline, rather than absolute neurotransmitter levels. Moreover, these fluctuations were correlated with behaviour, but since they are measured as relative changes, fibre photometry alone cannot establish a direct causal role of DA transients in decision-making and reward prediction. Nonetheless, numerous studies have demonstrated that NAc DA plays a crucial role in modulating behavioural outcomes of visual detection, attentional and discrimination tasks ^70,71,126–131^. In addition, ample research has demonstrated the role of the RPE in learning and decision-making. This study supports this function by demonstrating an active updating of the reward prediction in a manner consistent with a meta-cognitive interpretation in terms of confidence. However, there may be further roles for fluctuations in DA in the NAc as well as other striatal domains ^48,95,96,112,118,132,133^. In the present study, we did observe additional DA activity at trial initiation, perhaps related to the orienting response, although no systematic effects were identified. Moreover, several recent studies advocate an expanded RPE or more complex theory ^48,53,91–93,95,134–137^.

## Conclusion

Our findings illuminate novel functions of DA signalling in a visual signal detection task, highlighting its role in three distinct phases: trial initiation, decision-making and choice reward feedback. DA elevation across these phases aligns with the RPE theory, demonstrating that uncertainty introduced by behavioural manipulations and dynamic fluctuations in confidence continuously refine reward prediction and its error. Moreover, reward anticipation is dynamically updated in real-time, closely tied to decision-confidence and reflecting the complex interplay between uncertainty, confidence and reward processing. Notably, this intricate process occurred in response to both the presence and absence of a visual cue, with DA activity arising when a perceptual detection threshold was not met. These findings highlight the nuanced role of DA fluctuations in the NAcC, demonstrating their critical function in regulating decision-making processes. By linking real-time reward evaluation to confidence and uncertainty, this work underscores the neurochemical complexity underlying adaptive behaviour and provides a foundation for future studies exploring DA’s role in perceptual and executive functions.

## Methods

### Animals and housing

A total of 16 adult male Sprague Dawley rats (Envigo, Bichester, UK) housed in reversed-light cycle and housed in standard cages with wood-chip bedding and cage enrichment (a cardboard tunnel and woodblock) and kept in groups of four until surgeries, following an individual post-operative period the rats were pair-housed. Food restriction was applied when rats reached ∼250-300g bodyweight prior to behavioural experiments and restricted to 90% of their free-feeding bodyweight. There was *ad libitum* water available in the home cage, but not during behavioural experiments. This study has been conducted in accordance with the Animals (Scientific Procedures) Act 1986 Amendment Regulations 2012 (#PA9FBFA9F). Four rats were excluded from the final analysis, with one passing due to surgical complications and three exhibiting insufficient DA transients during the positive control test.

### Signal Detection task

The signal detection task (SDT) procedures are detailed by Turner and colleagues ^97–99^. Briefly, rats received training once or twice daily for 5-7 days a week. First, rats were trained to collect sugar pellets from both food magazines with a nose-poke in a magazine triggering the reward. Then, rats were trained to initiate trains by a nose-poke in the central port before collecting a pellet in one of the two food magazines. This was followed by signal detection in training stage 3, where rats had to learn to associate the illumination of a signal light with responding at one food magazine (signal trial) and the absence of the signal cue with responding at the other food magazine (no-signal trial) for a total of 120 trials with 60 signals trials and 60 no-signal trials. Once stable behaviour was achieved, variable signal durations (30, 60 and 250ms) were introduced to increase task difficulty for a minimum of 15 sessions to ensure stable individual performance, before commencing surgery. Which food magazine was assigned to signal trials was counterbalanced across the cohort but remained constant for each individual.

### Attentional Manipulations: Signal uncertainty and distraction

To increase the attentional load and introduce uncertainty, two behavioural manipulations were utilised; the short variable signal duration (short vSD) and visual distractor. The short vSD (10ms) tested the limit of individual visual perception, increasing task difficulty. The distractor manipulation utilized the house light. This manipulation had interleaved blocks (20 trials each) of standard and distractor trials, where the distractor would start blinking (1Hz) at the start of each trial until a response was recorded. Retaining the house light’s intended use of indicating a time-out period.

### Surgical procedures

Subjects were anaesthetised to surgically implant an optic fibre (Doric, Canada, 400µm core diameter 10mm length) at coordinates (AP +1.5, ML ±1.3, DV –6.3) and infuse the dLight1.3b viral vector (AAV1/2-hSyn1-chl-dLight1.3b-WPRE-bGHpA, Viral Vector Facility ETH Zurich v565-1, 2.5 × 10¹² GC/ml) at coordinates (AP +1.5, ML ±1.3, DV –6.1 & –6.5). Viral infusions were carefully administered using a Harvard Micro Syringe Pump Pico Plus and Hamilton 800 series 10µl syringes which were connected to a needle through plastic tubing (1µl virus was infused at 0.1µl/min for 10 minutes). Following each infusion, the needle was left in place for 5-10 minutes to allow for diffusion. Prior to the infusion and following the diffusion, the needle was briefly extended 0.05mm beyond the target depth to create a ‘pocket’ for the virus. This process was repeated for all coordinates. Subjects were placed in a stereotaxic frame using atraumatic bars whilst anaesthetised with isoflurane (5% for induction and 2-3% for maintenance), and the tooth bar was adjusted for a flat skull. Metal screws and dental cement were used to secure the optic fibre to the skull. The optic fibre implants were protected by a stainless – steel dust cap. Following surgery, rats were single housed and received oral Metacam (meloxicam oral suspension injected into soggy food pellets, Boehringer Ingelheim, Germany) for 3 days, before being pair housed and allowed to recover for the remaining days (≤7 days total).

### Fibre photometry recording

FP recordings were performed using Neurophotometrics FP3001 (Neurophotometrics, USA), which uses a blue 470nm LED and violet 415nm LED to excite the virally delivered fluorescent targets and detects light emission with a BlackFly camera (FP recordings were interleaved at 40Hz and 60mA). Prior to recording, the fibre optic cables were bleached at full power for a minimum of 30 minutes. Bonsai (Bonsai Foundation CIC, version 3002) was used to control the Neurophotometrics system, gather FP data and match behavioural events through external camera recording. The within Bonsai with camera recorded behavioural timestamps were then successfully matched to timestamps of behavioural events recorded with KLimbic (Med Associates) through a custom script in RStudio (RStudio Inc., version 2023.06.0, R v4.2.2) ^138^.

The Neurophotometrics system was connected to a fibre optic patch cord (BBP(2)_400/440/LWMJ-0.37_1m_FCM-2xFCM_LAF Branching bundle Patchcord – Low AutoFluorescence, Doric), which was attached to a rotary joint (FRJ_1×1_PT_400/440/LWMJ-0.37_1m_FCM_0.10m_FCM Fiberoptic Rotary Joint, pigtailed with 400 µm, NA0.37 fibre, 1 m input, 0.10 m output, optimized for FP, Doric) and subsequently to a fibre-optic patch cord (MFP_400/440/LWMJ-0.37_0.65m_FC-CM3_LAF Mono Fiberoptic Patchcord – Low AutoFluorescence, Doric) that was attached to the 400 µm fibre optic implant.

Prior to connecting the subjects to the cable both lasers were extinguished to avoid blinding the subjects. The patch cord was then securely attached to the fibre optic implant and the rat was placed in the operant chamber. The FP recording was started in Bonsai and subsequently the 415 nm isosbestic channel was turned on and after a pause of several seconds the 470 nm signal channel was turned on. The onset of the traces was visually inspected. The rat was then presented with several (∼3) sucrose pellets to briefly re-validate positive DA signals. Finally, the operant chambers were closed, and the behavioural test was started in the KLimbic programme. All Bonsai inputs and behavioural results were monitored to ensure everything was running smoothly. When necessary, the patch cord was tightened during the behavioural task. Once the task finished, the rats were again presented with sucrose pellets to re-validate positive DA-signals. The 470 nm signal was then turned off and finally the 415 nm isosbestic channel was turned off. The Bonsai recording was stopped, and the rat was detached from the patch-cord.

### Histology

Following the conclusion of behavioural testing, brains were harvested following transcardial perfusion with 0.01M PBS followed and 10% formalin saline. Brains were submersed in 20% ^w^/_v_ sucrose (Fisher chemical, S/8560/63) after 24 hours. Brains were sliced at 60mm on a freezing microtome and stored in cryoprotectant solution at –20°C. Cryoprotectant solution was prepared with 30% ^w^/_v_ sucrose, 30% ^v^/_v_ ethylene glycol (Acros Organics, 444230010), 0.547% ^w^/_v_ disodium hydrogen orthophosphate, 0.159% ^w^/_v_ sodium dihydrogen orthophosphate (Signal-Aldrich, 71500) and 0.9% ^w^/_v_ sodium chloride in deionised water at 60°C, following the dissolution of the above mentioned components the 1% ^w^/_v_ polyvinyl pyrrolidone (Acros Organics, 227541000) was added.

Free-floating brain slices were washed 3×10 min in PBS and blocked (1h at RT on shaker) in blocking buffer (BB) (0.01M PBS, 1% w/v Bovine serum albumin (Sigma-Aldrich, A7906) and 0.3% v/v Triton X-100 (Sigma-Aldrich, 3787). Followed by incubation (overnight at RT on shaker) with chicken anti-GFP (1:1000, ab13970 Abcam) and rabbit anti-GFAP (1:1000, A-85419 Antibodies.com) in BB. Slices were washed 3x 10min with PBS and incubated (2h at RT in the dark) with goat anti-chicken (1:2000, ab175477 Abcam, AlexaFluor568) and goat anti-rabbit (1:2000, A-11008 ThermoFisher Scientific, AlexaFluor488) in BB. Finally, after 3x 10min PBS wash, brain slices were mounted on Menzel Gläzer Superfrost® Plus (Thermo Fisher Scientific, J1800AMNZ) using FluorSave™ Reagent (Calbiochem, 345789). Immunohistochemistry visualization and quantification was carried out using a fluorescent microscope (Zeiss AXIO Imager M2 set to 20x magnification) and the software programme Visiopharm.

### Preprocessing of fibre photometric data

The 470nm and 415nm, dLight signal and isosbestic control excitation wavelengths were recorded by the system. The sole integrated preprocessing step from the system was the data scaling. All other processing steps were performed individually by subject and session in Spyder (Spyder IDE, version 5.4.3 ^139^, Python 3.11.5 ^140^) through custom scripts. First, data were loaded and correctly labelled isosbestic and signal. Data were cropped by subject to only include datapoints from the behavioural session as well as datapoints from the 5s preceding the start of the session and to the 5s after the completion of the SDT session. Data were then filtered with a zero-phase moving average filter of 1 s in accordance with the method of Sherathiya and colleagues (2021) ^141^. The isosbestic control was fitted to the signal as linear decay using least square linear regression. Then the delta F/F and subsequent z-scores were calculated over the whole fitted signal trace and Peri-stimulus/event time histograms (PSTH) of 6s were extracted to examine responses to discrete behavioural events over repeated trials. The individual datapoints within each PSTH were re-baselined by means of subtraction of the mean z-score of 5 baseline trials, inclusive of the current trial.

### Frequency filtering

Offline fibre photometry raw data were processed using custom Python 3.11.5 scripts using scipy package ^142^ to analyse activity fluctuations across different frequency bands over time. Fiber photometry data were filtered into four different frequency bands using a 4^th^-order Butterworth bandpass filter at 0.15 to 0.6 Hz, 0.6 to 2.5 Hz and 2.5 to 10 Hz, as well as a 4^th^–order Butterworth lowpass filter with a cutoff frequency of 0.15 Hz. The rationale behind these bands was that Jørgensen and colleagues (2023) reported slower and faster dynamics on DA-dependent ventral and dorsal striatum, respectively ^100^. This processing was done using a forward-backward filtering to avoid the potential frequency-dependent phase shifts ^143^. Subsequently, data were segmented into epochs of 6 s (–3 to 3 s) with the 0 s centred on the response.

### Behavioural analysis

Two-choice tasks commonly report accuracy, however analysis in accordance with the signal detection theory is more appropriate ^144–146^. First, the number of Hits, Misses, False Alarms (FAs) and Correct Rejections (CRs) were calculated. A hit was defined as correctly reporting the presence of the visual cue, while a CR was the correct response to the absence of the visual cue. A miss occurred when there was a visual cue and the rat reported that there was no cue, while the FA was the incorrect detection of the visual cue. Thus, both correct responses were rewarded, while their incorrect counterparts prompted a 5s time-out. Subsequently, the sensitivity and bias were calculated according to the signal detection theory. Sensitivity refers to the capacity of a subject to distinguish between signal and noise trials and was approximated with d’ (**Equation 1**). Bias refers to the subject’s preference for reporting signal or noise (no-signal trials) and was approximated by β (**Equation 2**).

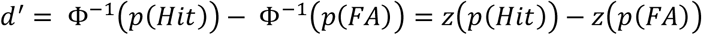

**Equation 1**: Calculation of sensitivity d’, where φ^-1^ = z and returns a z-score. p(Hit) and p(FA) indicate the probability of making a hit and false alarm, respectively. The d’ is in standard deviation units, where 0 indicates a complete inability to distinguish signal from noise, and larger positive values indicate an increasingly greater ability to distinguish signals from noise. Negative values arise through sampling error or response confusion.

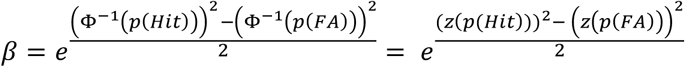

**Equation 2**: Calculation of bias β using the standardized probability of making a hit and false alarm. A β value of 1 signifies no bias, β < 1 a bias towards the reporting the signal and β > 1 a bias towards reporting its absence.

### Data analysis and visualisation

Following histological analysis (see Fig. 6) and the positive FP control tests, a total of n = 12 rats was included in the study. Baseline DA transients were recorded across three consecutive baseline SDT sessions. Transients were averaged by session per trial type before averaging by subject. Subsequently, these values were averaged and used to obtain a representative DA transient for each trial type. To assess differences in DA transients, the signal detection theory trial division was utilised, resulting in distinct DA transients for Hits, Misses, FAs and CRs. Additionally, DA transients were grouped according to the signal duration (no-signal, 30ms, 60ms, 250ms and 1s). Due to the rapid succession of events within a trial, the results could only be accurately aligned with a singular event. Data were aligned to trial initiation, which coincided with the onset or absence of the signal light, as well as to the recorded nose-poke decision, coinciding with the reward delivery or onset of the house light.

**Figure 6:**
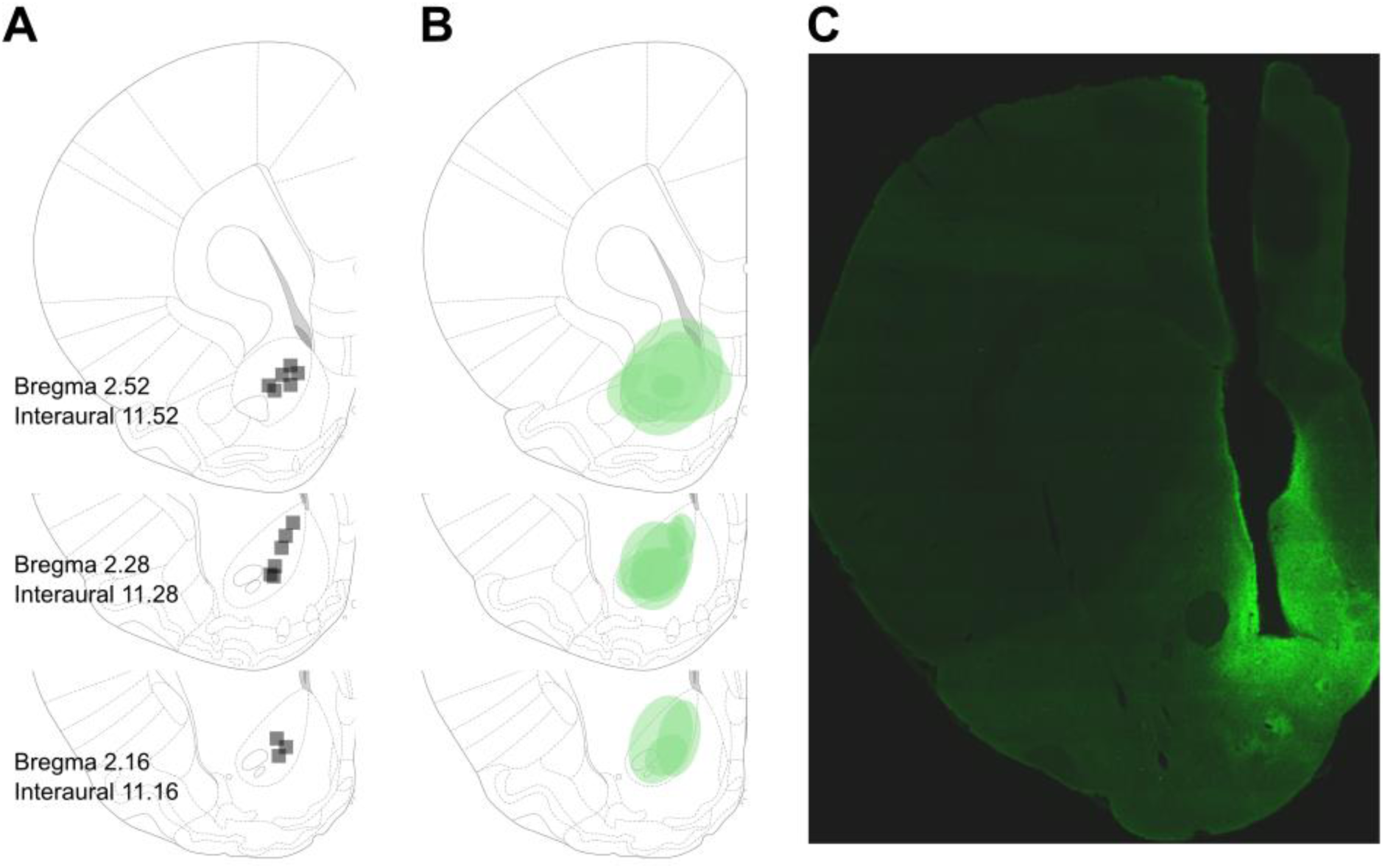
Site of the fibre optic placement and viral expression of dLight1.3b in the nucleus accumbens core by subject. (A) Fibre optic placement. (B) Viral expression of GFP attached to dLight1.3b. (C) Representative example of GFP expression.

The trial period was divided into three timeframes of interest: (1) trial initiation; (2) decision –making period (≥1 second); and (3) feedback period (Fig. 1A). Due to the complexity of the SDT resulting in probable differences in DA transients, various distinctions in trial type were made based on outcome categories, outcome dependent on the previous outcome and signal duration. DA transients were compared both visually (qualitatively) and quantitatively through the area under the curve (AUC) values with mixed-model ANOVAs with restricted maximum likelihood. The ANOVAs included the following variables: behavioural outcome (within-subject), time frame (within-subject), distractor (within-subject, optional), response speed (within-subject, optional), the rat ID was a random effect to accommodate for repeated measures. Post-hoc comparisons were performed where appropriate and the degrees-of-freedom methods used was Kenward-Roger, Tukey’s method was used for p-value adjustment.

## Acknowledgements

We acknowledge Shionogi & Co., Ltd for funding this research. We thank dr. B.J. Alsiö for helpful discussions and foundational work, Oliwia Stecko and Merak Chen for their experimental assistance, dr. Aske Ejdrup for his support during the fibre photometry preprocessing pipeline, and Dr. Armin Lak for his insights into confidence signals in the nucleus accumbens. The experimental work was carried out under a Home Office Project Licence held by Dr. A. L. Milton.

## Author contributions

LJFWV designed and conducted experiments, collected data, wrote data analysis scripts, analysed data and wrote the manuscript. CM designed and conducted experiments, collected data, provided technical support and wrote the manuscript. FEA wrote data analysis scripts and wrote the manuscript. KY designed experiments and wrote the manuscript. JWD designed experiments and wrote the manuscript. TWR secured funding, designed experiments and wrote the manuscript.

