## Supplementary data for "Dopamine Dynamics in the Nucleus Accumbens Reflect Confidence in Detecting the Occurrence and Non-Occurrence of Visual Signals in Perceptual Decision-Making"

### Supplementary figures

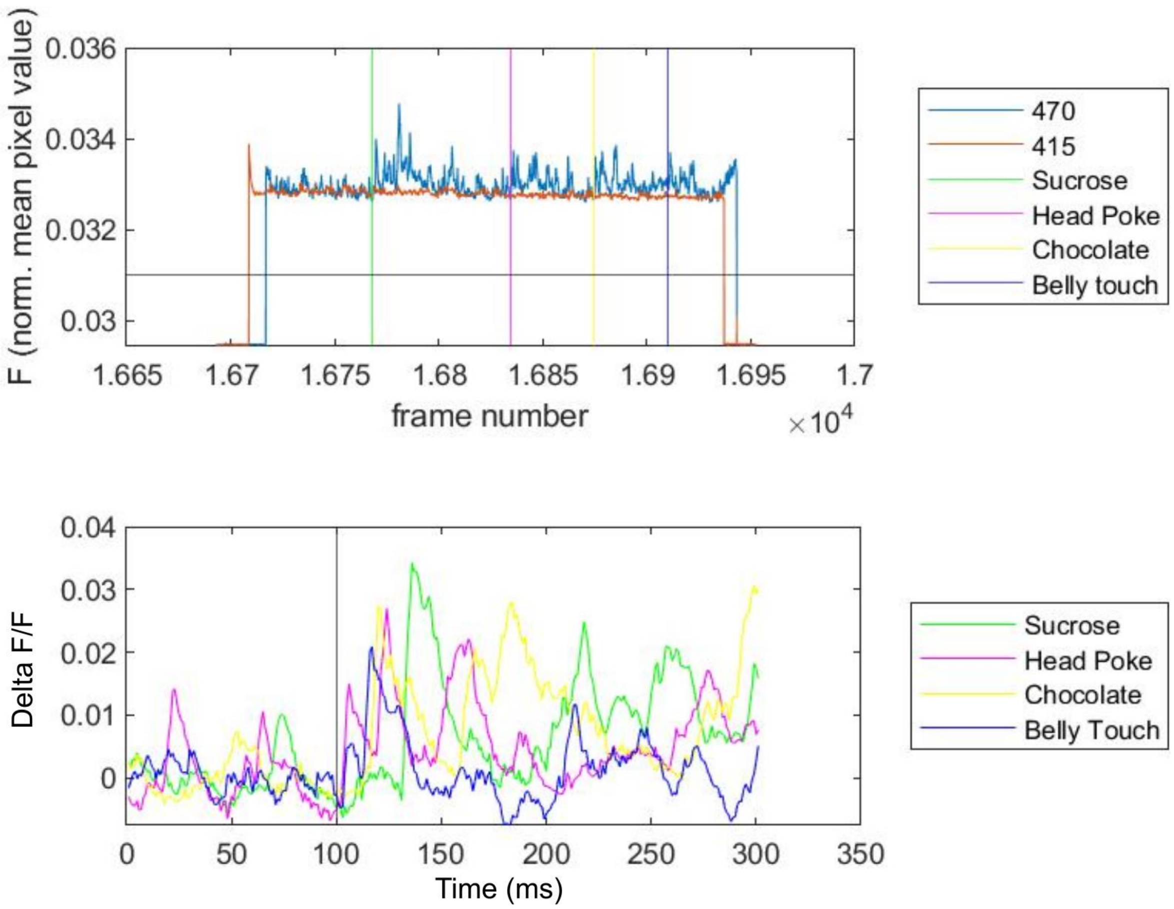


S1: Representative unprocessed raw data and sucrose test results.

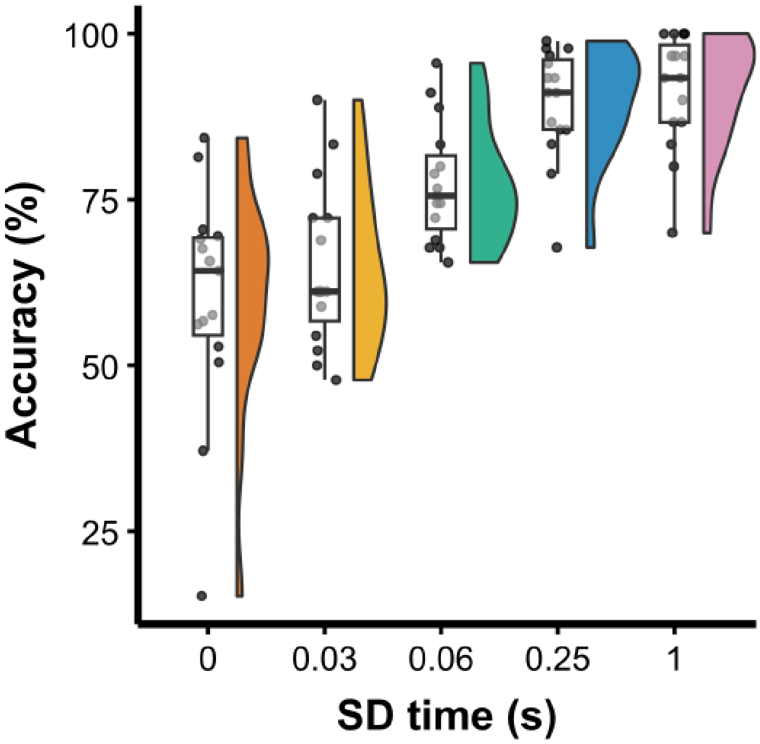


S2: Signal detection performance (Accuracy) for different signal durations (n = 15). Average accuracy by signal duration across three baseline sessions by subject.

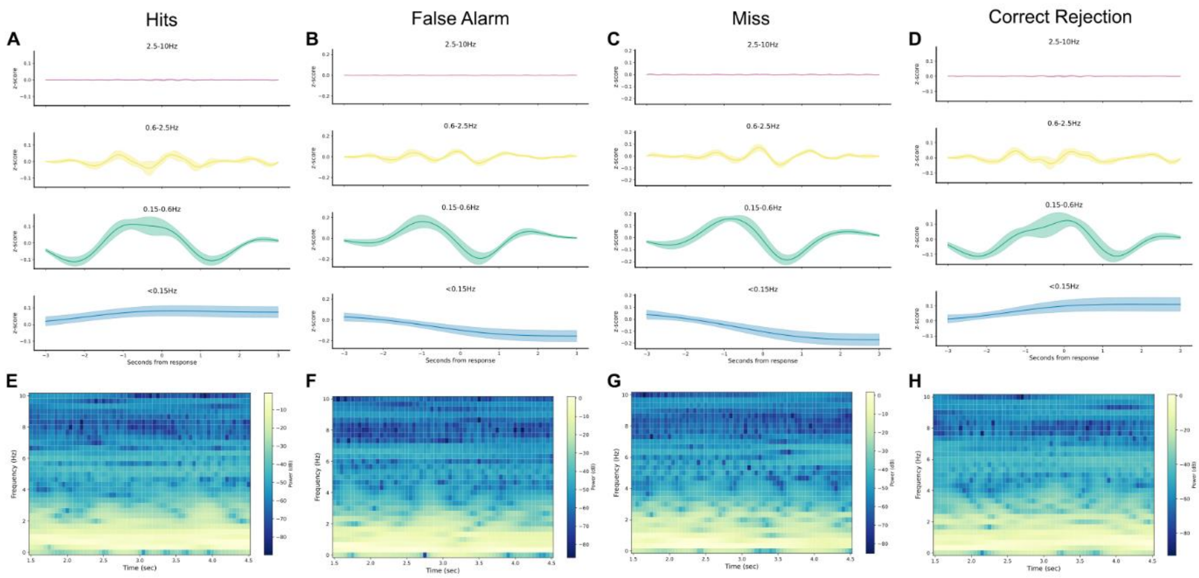


S3: Filtered fibre photometry traces by signal detection theory outcomes and a representative spectrogram. (A-D) Average DA transients by frequency range across three baseline sessions by subject. (E-H) Representative spectrograms of a trial for each signal detection theory outcomes.

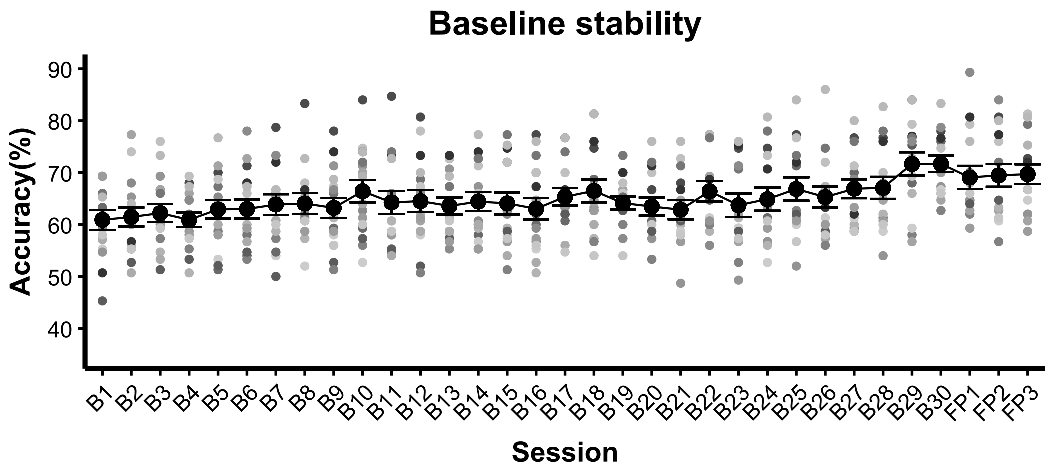


S4: Signal detection performance (Accuracy) across sessions.

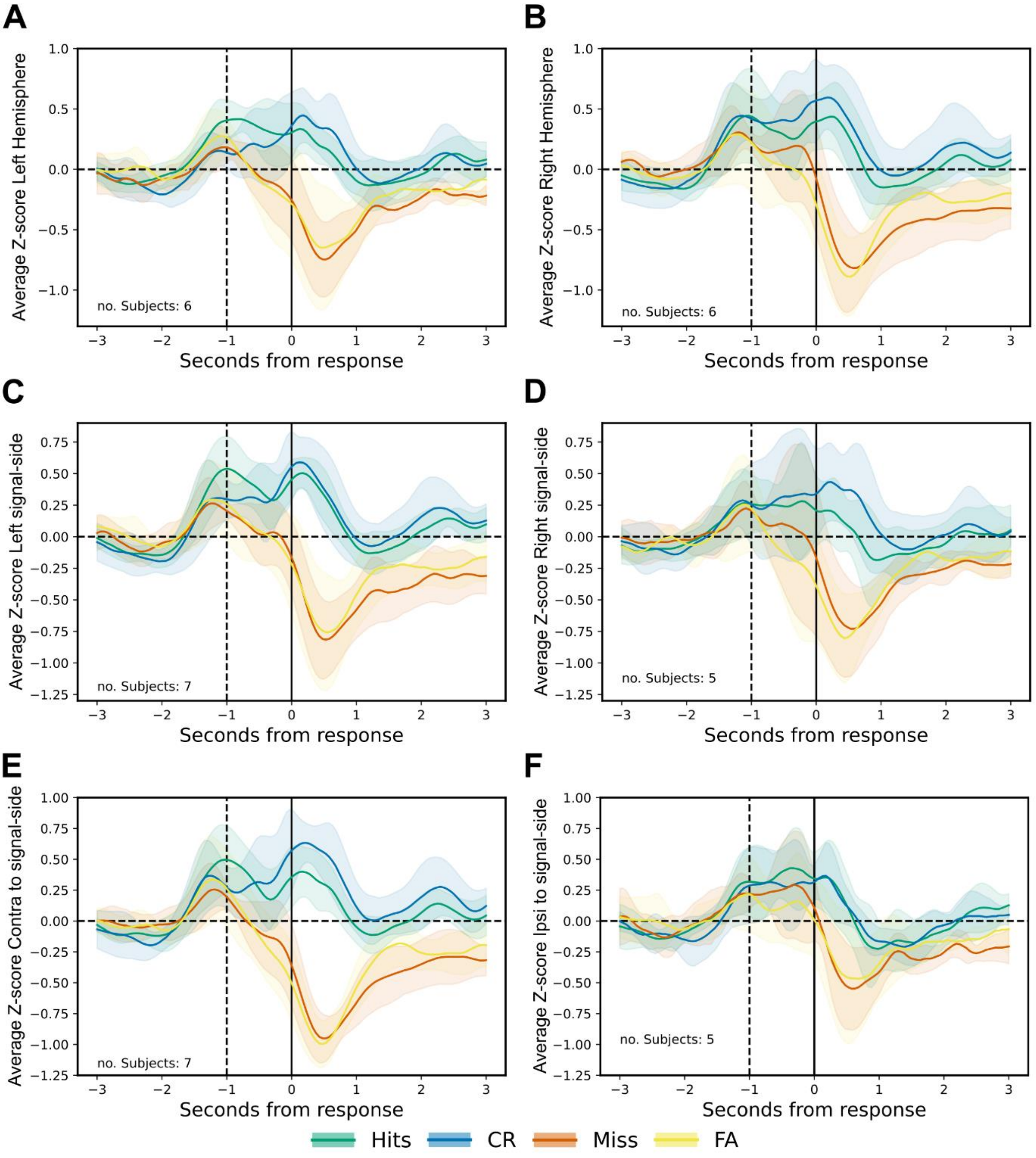


S5: DA transients measured during SDT performance for each signal detection theory outcome. Results were averaged across three sessions and depict the average Z-score ± 95% confidence interval. Subjects with fibre optic implanted in left hemisphere (A) and right hemisphere (B). Subjects which had to respond to the left (C) and right (D) food magazine to correctly identify the signal cue. Subjects with optic fibre implanted in the hemisphere contralateral (E) and ipsilateral (F) to the food magazine side associated with correct identification of the signal cue.
